# *In vitro* and *in vivo* venom-inhibition assays identify metalloproteinase-inhibiting drugs as potential treatments for snakebite envenoming by *Dispholidus typus*

**DOI:** 10.1101/2022.01.07.475313

**Authors:** Stefanie K. Menzies, Rachel H. Clare, Chunfang Xie, Adam Westhorpe, Steven R. Hall, Rebecca J. Edge, Jaffer Alsolaiss, Edouard Crittenden, Robert A Harrison, Jeroen Kool, Nicholas R. Casewell

## Abstract

Snakebite envenoming affects more than 250,000 people annually in sub-Saharan Africa. Envenoming by *Dispholidus typus* (boomslang) results in venom induced consumption coagulopathy, whereby highly abundant prothrombin-activating snake venom metalloproteinases (SVMPs) consume clotting factors and deplete fibrinogen. The only available treatment for *D. typus* envenoming is the monovalent SAIMR Boomslang antivenom. Treatment options are urgently required because this antivenom is often difficult to source and, at $6,000/vial, typically unaffordable for most snakebite patients. We therefore investigated the *in vitro* and *in vivo* preclinical efficacy of four SVMP inhibitors to neutralise the effects of *D. typus* venom; the matrix metalloproteinase inhibitors marimastat and prinomastat, and the metal chelators dimercaprol and DMPS. The venom of *D. typus* exhibited an SVMP-driven procoagulant phenotype *in vitro*. Marimastat and prinomastat demonstrated equipotent inhibition of the SVMP-mediated procoagulant activity of the venom *in vitro*, whereas dimercaprol and DMPS showed considerably lower potency. However, when tested in preclinical murine models of envenomation, DMPS and marimastat demonstrated partial protection against venom lethality, demonstrated by prolonged survival times of experimental animals, whereas dimercaprol and prinomastat failed to confer any protection at the doses tested. The results presented here demonstrate that DMPS and marimastat show potential as novel small molecule-based therapeutics for *D. typus* snakebite envenomation. These two drugs have been previously shown to be effective against *Echis ocellatus* venom induced consumption coagulopathy (VICC) in preclinical models, and thus we conclude that marimastat and DMPS may be valuable early intervention therapeutics to broadly treat VICC following snakebite envenoming in sub-Saharan Africa.

## 1. Introduction

More than 250,000 cases of snakebite envenoming are estimated to occur annually in sub-Saharan Africa^1^, disproportionately affecting those in rural, impoverished communities without adequate access to healthcare^2,3^. Venom-induced consumption coagulopathy (VICC) is a common manifestation of snakebite envenoming, during which procoagulant venom toxins consume clotting factors resulting in the ensuing depletion of fibrinogen and, ultimately, coagulopathy^4^. Several clotting factors are the target for procoagulant snake venom toxins, and these include Factor X, Factor V, fibrinogen and prothrombin^4^. While infrequent, envenomings by the rear fanged African colubrid *Dispholidus typus* (boomslang) are characterised by causing VICC^5–7^. The venom of this species is known to potently activate prothrombin^8^, resulting in the liberation of thrombin, and the subsequent downstream consumption of fibrinogen and fibrin, causing dysregulation of coagulation^8–10^. The activation of prothrombin is likely the result of snake venom metalloproteinases (SVMPs)^9,10^, which are the dominant toxin type present in the venom and account for almost 75% of the proteinaceous toxins^11,12^. Other minor toxin families identified in the *D. typus* venom proteome include three-finger toxins, phospholipases A2 (PLA_2_s), cysteine-rich secretory proteins (CRISPs), snake venom serine proteases and C-type lectin-like toxins (of which each constitute <10% of the venom proteome)^11^, though their contribution to envenoming pathology remains unclear.

*Dispholidus typus* is broadly distributed throughout much of sub-Saharan Africa (sSA) and whilst incidences of envenoming are fortunately rare, the rapid and severe VICC consequences pose considerable clinical challenges. This is because the only specific treatment for *D. typus* envenoming is the monospecific F(ab’)_2_ antivenom “SAIMR Boomslang” (South African Vaccine Producers Pty Ltd), which has limited availability outside of the Southern Africa Economic Community, and costs as much as US$6050 per vial^13^. Given that *D. typus* exhibits a broad geographical distribution throughout much of sub-Saharan Africa (sSA), the only specific for treatment for envenomings caused by this species is largely unobtainable for snakebite victims who either cannot afford, or do not have access to, the antivenom^9^, and thus investigating novel treatments is a research priority.

More generally, it is well recognised that conventional polyclonal antibody-based antivenoms have several shortcomings, despite being life-saving therapeutics. In addition to often being unaffordable to many snakebite victims, they are associated with high rates of adverse reactions^14,15^, and have poor dose efficacy, with only ~10-20% of the active immunoglobulins recognising and binding to venom toxins^16^. Logistically, antivenoms are poorly suited for the rural locations in which they are typically required; for example many antivenoms rely on cold chain transport and storage and must be administered intravenously by trained staff in healthcare facilities^17^. Indeed, up to 75% of deaths from snakebite are estimated to occur before patients are able to reach healthcare facilities, thus there is a compelling need to identify novel snakebite treatments that could be administered in the community soon after a bite^18^.

To this end, small molecule-based drugs (i.e. ‘toxin inhibitors’) have received considerable interest as novel snakebite therapeutics, both as potential individual treatments or in combination with existing antivenoms^18–20^. Small molecule drugs have a number of potentially advantageous characteristics over antivenoms, including improved affordability and stability, oral delivery format, higher tolerability^20^, and improved tissue penetration^21,22^. Previously, Ainsworth *et al* demonstrated the *in vitro* inhibitory effect of the metal chelator EDTA against SVMP-mediated prothrombin degradation caused by *D. typus* venom, suggesting that small molecule inhibitors may be effective therapeutics for *D. typus* envenoming^9^. In the same study, Ainsworth *et al* demonstrated in a murine preclinical model that EDTA was protective against the lethal effects of *Echis ocellatus* venom, an African viper which, similar to *D. typus* venom, contains a high abundance of SVMP toxins^9^, including prothrombin activators, and causes VICC in envenomed victims^23,24^. Other small molecule drugs with SVMP-inhibiting potential include other metal chelators, such as dimercaprol and DMPS (2,3-dimercapto-1-propanesulfonic acid)^25–27^, and the mimetic matrix metalloprotease inhibitors marimastat, batimastat and prinomastat^22,24,28^. Marimastat and batimastat were found to effectively inhibit SVMP activity and reduce haemorrhagic pathologies in murine models of *E. ocellatus* envenoming^24^, and inhibit the procoagulant effects of several viper venoms *in vitro*^24,25^. Similarly, prinomastat (AG-3340) showed inhibitory activity against the haemorrhagic effects of both purified SVMPs and the crude venom of *E. ocellatus*^28^. The metal chelators dimercaprol and DMPS have also been shown to inhibit SVMP activity of *E. ocellatus* venom *in vitro*, with DMPS also demonstrating *in vivo* preclinical neutralisation of venom lethality and haemorrhage^26^. While *D. typus* venom is dominated by SVMPs, PLA_2_ toxins are also thought to contribute to the coagulopathy induced by this venom^29^. The small molecule drug varespladib has been extensively investigated as an inhibitor of venom PLA_2_ toxins found in a wide geographical range of snake species, with such studies showing potent neutralisation of PLA_2_ activity^25,27,30^ and associated anticoagulant, haemorrhagic, myotoxic and neurotoxic pathologies^31–35^. Thus, small molecule treatments for snakebite have the potential to overcome the current species-specific and restrictive geographical utility of current antivenoms.

Despite these promising recent research outcomes, further investigation is required to explore the inhibitory breadth and potency of small molecule toxin inhibitors due to the ubiquitous variability in snake venom composition and therefore, also, the variant toxin specificities of these inhibitory small molecule drugs. In particular, despite overarching similarities in venom composition and ensuing snakebite pathology between *Echis* spp. and *D. typus*^9^, the SVMPs of *D. typus* have evolved their prothrombin activating ability independently of those found in the venom of *Echis* spp.^10^. Consequently, building on previous principles demonstrated for *Echis* spp.^24–26,36^, in this study four small molecule drugs were investigated *in vitro* and *in vivo* to assess their inhibitory potential against the venom of *D. typus*. To do so, we applied *in vitro* metalloproteinase and coagulation bioassays on crude and nanofractionated venom, and *in vivo* murine models of envenoming to assess neutralisation of venom lethality.

## 2. Methods

### 2.1 Venoms

Lyophilised *D. typus* venom (Product code L1403, origin South Africa, purity >99%) was sourced from Latoxan (Portes les Valence, France) and stored at 4 °C to ensure long-term stability. Prior to use, venom was resuspended in PBS (pH 7.4, Gibco) at 1 mg/mL for *in vitro* experiments and 5 mg/mL for *in vivo* experiments.

### 2.2 Drug preparations for *in vitro* studies against crude venom

The small molecule SVMP inhibitors tested were; dimercaprol (2,3-dimercapto-1-propanol, ≥98 % iodometric, Cat no: 64046, Sigma), DMPS (2,3-dimercapto-1-propane-sulfonic acid sodium salt monohydrate, 98%, Cat no: H56578, Alfa Aesar), marimastat ((2*S*,3*R*)-*N*4-[(1*S*)-2,2-Dimethyl-1-[(methylamino)carbonyl]propyl]-*N*1,2-dihydroxy-3-(2-methylpropyl)butanediamide, >98%, Cat no: 2631, Tocris Bioscience), prinomastat hydrochloride (Cat no: HY-12170A, >98%, MedChemExpress). Varespladib (2-[[3-(2-Amino-2-oxoacetyl)-2-ethyl-1-(phenylmethyl)-1H-indol-4-yl]oxy]-acetic acid, Cat no: SML1100, >98% HPLC, Sigma) was used as a small molecule drug control. All drugs were reconstituted in dimethyl sulfoxide (DMSO) (Sigma) to 10 mM stocks and stored at −20 °C. Daughter plates were created at 1 mM concentrations in 384-well format to allow the creation of assay-ready plates using a VIAFLO 384 electronic pipette (Integra). Both daughter plates and assay-ready plates were stored at −20 °C, with the latter used within a month of creation. For the SVMP assay 0.91 μL of each drug was plated (final reaction volume of 91 μL), while 0.5 μL was plated for the coagulation assay (final reaction volume of 50 μL). For marimastat, prinomastat and varespladib, dose response curves were created at a final concentration range of 10 μM to 4.8 pM using a two-fold dilution (50 μL drug into 50 μL of DMSO), with each concentration tested in duplicate. For DMPS and dimercarpol, dose response curves were created at a final concentration range of 160 μM to 76.3 pM using a two-fold dilution (50 μL drug into 50 μL of DMSO), with each concentration tested in duplicate.

### 2.3 *In vitro* neutralisation of coagulopathic crude venom activity

To assess the inhibitory potency of the selected drugs against coagulopathic venom activity we used a previously described absorbance-based plasma clotting assay^37^. Citrated bovine plasma (VWR) was defrosted and centrifuged at 858 x g for 5 minutes to remove precipitates before use. Thereafter, 100 ng of venom in 10 μL PBS was added to each well in the 384-well assay-ready plate (containing 0.5 μL of 1 mM of inhibitor) using a VIAFLO 384, the plate was then briefly spun down in a Platefuge (Benchmark Scientific) and incubated at 37 °C for 25 minutes, followed by a further five minutes acclimatisation at room temperate. Next, 20 μL of 20 mM CaCl_2_ was added using a MultiDrop 384 Reagent Dispenser (ThermoFisher Scientific), followed by the immediate addition of 20 μL citrated bovine plasma. The plate was then immediately read for kinetic absorbance at 595 nm for 116 minutes using a FLUOstar Omega platereader (BMG Labtech).

Assays were performed in triplicate and each assay contained technical duplicates at each dose. Positive control values were generated using DMSO + venom, and negative control values were generated using DMSO in the absence of venom. All compounds were analysed for their ability to return clotting to normal at the timepoint at which the positive and negative absorbance values were furthest apart. For this, the raw values were normalised to show percentage of normal clotting, e.g. a value of 100% meant the compound returned clotting to that of the negative control. These percentage values were plotted and fitted with a nonlinear regression curve for the normalised response (variable slope) using to calculate the IC_50_ data and 95% confidence intervals for each compound using GraphPad Prism 9.0 (GraphPad Software, San Diego, USA). Multiple comparisons one-way ANOVA test was used to compare IC_50_ values generated for each replicate plate, using GraphPad Prism 9.0.

### 2.4 Venom nanofractionation

To further explore the inhibitory specificity of the selected drugs, we fractionated *D. typus* into toxin constituents and repeated the plasma bioassay. Venom nanofractionation^29,38^ was performed on a UPLC system (‘s Hertogenbosch, The Netherlands) controlled by Shimadzu Lab Solutions software. Venom solution was prepared by dissolving lyophilised *D. typus* venom into water (purified by Milli-Q Plus system, Millipore) to a concentration of 5.0 mg/mL and stored at −80 °C until use. For each analysis, 50 μL venom solution (1.0 mg/mL) was injected by a Shimadzu SIL-30AC autosampler after diluting the stock venom solutions (5.0 ± 0.1 mg/mL) in Milli-Q water. A Waters XBridge reversed-phase C18 column (4.6 × 100 mm column with a 5 μm particle size and a 300 Å pore size) was used for gradient separation at 30 °C. Mobile phase A was composed of 98% water, 2% acetonitrile (ACN) (Biosolve) and 0.1% formic acid (FA) (Biosolve), while mobile phase B was composed of 98% ACN, 2% water and 0.1% FA. The total solvent flow rate was maintained at 0.5 mL/min and the gradients were run as follows: linear increase of eluent B from 0 to 50% in 20 min followed by a linear increase to 90% B in 4 min, then isocratic elution at 90% for 5 min, subsequently the eluent B was decreased from 90% to 0% in 1 min followed by an equilibration of 10 min at 0% B. The column effluent was split as two parts (9:1), with the smaller fraction (10%) sent to a Shimadzu SPD-M20A prominence diode array detector. The larger fraction (90%) was directed to a FractioMate nanofractionator (SPARK-Holland & VU) and fractions collected onto transparent 384-well plates (F-bottom, rounded square well, polystyrene, without lid, clear, non-sterile; Greiner Bio One). The nanofractionator was controlled by FractioMator software (Spark-Holland) to collect fractions continuously at a resolution of 6 s/well. After collection, the well plates with venom fractions were dried overnight in a Christ Rotational Vacuum Concentrator (RVC 2-33 CD plus, Zalm en Kipp, Breukelen, The Netherlands), to remove any solvent remaining in the wells. The vacuum concentrator was equipped with a cooling trap maintained at −80 °C during operation. The dried plates were then stored at −20 °C until bioassaying.

### 2.5 *In vitro* neutralisation of coagulopathic venom toxin fractions

The small molecule inhibitors marimastat ((2S,3R)-N4-[(1S)-2,2-Dimethyl-1-[(methylamino)carbonyl] propyl]-N1,2-dihydroxy-3-(2-methylpropyl) butanedia-mide), prinomastat hydrochloride (AG-3340 hydrochloride), dimercaprol (2,3-Dimercapto-1-propanol), DMPS (2,3-dimercapto-1-propane-sulfonic acid sodium salt monohydrate) and varespladib (A-001) were purchased from Sigma-Aldrich. Bovine plasma (Sodium Citrated, Sterile Filtered, Product Code: S0260) was purchased from Biowest. For assay preparation, the CaCl_2_ (Biosolve), which was used to de-citrate plasma to initiate coagulation in the coagulation assay, was dissolved in Milli-Q water to 20 mM. The inhibitors were dissolved in DMSO (≥ 99.9%, Sigma-Aldrich) to a concentration of 10 mM and stored at −20 °C. The plasma was defrosted and then centrifuged at 2000 rpm (805 × g) for 4 min in a 5810 R centrifuge (Eppendorf) to remove possible particulate matter. The inhibitor stock solutions were diluted in PBS buffer to the described concentrations, then 10 μL of each diluted inhibitor solution was pipetted to all wells of plate wells containing freeze-dried nanofractionated venom fractions by a VWR Multichannel Electronic Pipet, followed by centrifuging the plate for 1 min at 805 x g. Next, a pre-incubation step for 30 min at room temperature was performed. Final concentrations of inhibitor solutions used for the coagulation bioassay were 20, 4, 0.8, 0.16 and/or 0.032 μM (with corresponding DMSO final concentrations of 0.02%, 0.004%, 0.0008%, 0.00016% and 0.000032%, respectively). After this incubation step, the HTS coagulation assay was performed as described by Still *et al* ^37^. A Multidrop 384 Reagent Dispenser (Thermo Fisher Scientific) was used to dispense 20 μL of CaCl_2_ solution onto all wells of the 384-well plates, followed by 20 μL plasma after rinsing of the Multidrop with deionized water between dispensing. Kinetic absorbance measurements were conducted immediately for 100 min at 595 nm and 25 °C using a Varioskan Flash Multimode Reader (Thermo Fisher Scientific). Venom-only analyses were performed as control experiments, for which 10 μL PBS instead of inhibitor solution was added to all wells of the vacuum-centrifuge-dried nanofractionated well plates. Each nanofractionation analysis was performed in at least duplicate.

The resulting coagulation chromatograms were plotted as described by Slagboom *et al*^29^, with each chromatogram reconstructed to display ‘very fast coagulation’, ‘slightly/medium increased coagulation’ and ‘anticoagulation’. To plot the very fast coagulation chromatogram, the average slope of the first five minutes in the assay was plotted, and for the slightly/medium coagulation chromatogram the average slope of the first 20 minutes was plotted. For anticoagulant chromatogram the final (end-point) read at 100 minutes was plotted. Clotting velocities were all plotted against the venom nanofractionation time, producing positive peaks for procoagulant compounds and negative peaks for anticoagulant compounds.

### 2.6 *In vitro* neutralisation of venom SVMP activity

The SVMP activity of crude *D. typus* venom in the presence of inhibitors or vehicle control (DMSO), was measured using a quenched fluorogenic substrate (ES010, R&D Biosystems), in line with principles previously described^26^. The substrate was suspended in reaction buffer (150 mM NaCl, 50 mM Tris-HCl pH 7.5) and used at a final concentration of 10 μM (supplied as a 6.2 mM stock). Reactions consisted of 1 μg of venom (1 μg in 15 μL PBS) co-incubated with 0.91 μL of 1 mM of inhibitor. The 384 well plate (Greiner) was briefly spun down in a Platefuge (Benchmark Scientific) and incubated at 37 °C for 25 minutes, with an additional 5 minutes acclimatisation at room temperate, before the final addition of the freshly diluted fluorogenic substrate (75 μL of 12.1 μM). The plate was immediately run on a CLARIOstar platereader (BMG Labtech) at an excitation wavelength of 320-10 nm and emission wavelength of 420-10 nm with 10 flashes per well at 25 °C for 100 cycles (each cycle time 79 seconds). The assay was performed independently in technical duplicate. The end-reads were calculated for each sample at the time where all fluorescence curves had typically reached a plateau (maximum fluorescence). SVMP activity was calculated for each test condition as a percentage of the mean of the DMSO only wells (100% activity), with a baseline of the marimastat 10 μM controls representing 0% activity. IC_50_ values were calculated from the percentage inhibition values by fitting a nonlinear regression curve for the normalised response (variable slope) for each compound using GraphPad Prism 9.0 (GraphPad Software, San Diego, USA). The best-fit IC_50_ values for each replicate were compared to identify significant differences between the IC_50_ values for each drug using one-way multiple comparisons ANOVA analysis in GraphPad Prism 9.0.

### 2.7 *In vivo* neutralisation of venom lethality

#### 2.7.1 Animal ethics

All animal experiments were performed using protocols approved by the Animal Welfare and Ethical Review Boards of the Liverpool School of Tropical Medicine and the University of Liverpool, under project licence (P58464F90) approved by the UK Home Office in accordance with the UK Animal (Scientific Procedures) Act 1986.

#### 2.7.2 Animal maintenance

Male CD1 mice (18-20g) were sourced from Charles River (UK) and acclimatised for a minimum of one week before experimentation. Mice were grouped in cages of five, with room conditions of approximately 22 °C at 40-50% humidity, with 12/12 hour light cycles, and given *ad lib* access to CRM irradiated food (Special Diet Services, UK) and reverse osmosis water in an automatic water system. Mice were housed in specific-pathogen free facilities in Techniplast GM500 cages containing Lignocell bedding (JRS, Germany), Sizzlenest zigzag fibres as nesting material (RAJA), and supplied with environmental enrichment materials.

#### 2.7.3 Co-incubation model of preclinical efficacy

The median murine lethal dose (LD_50_) for *D. typus* venom administered by intravenous injection was previously determined as 22.29 μg per mouse^9^. To determine the efficacy of small molecule inhibitors against *D. typus* venom, a refined version of the WHO recommended antivenom ED_50_ neutralisation experiments was used, in which ~4 x LD_50_ doses of venom (90 μg) were pre-incubated with each small molecule inhibitor. Drug stocks were freshly prepared to allow for a ratio of 1:1.33 venom to inhibitor as previously defined by marimastat *in vivo* testing against other snake venoms^36^. Drugs tested *in vivo* were dimercaprol (2,3-dimercapto-1-propanol ≥98 % iodometric, Cat no: 64046, Sigma-Aldrich), marimastat (>98% HPLC, Cat no: M2699, Sigma-Aldrich), and prinomastat hydrochloride (≥95% HPLC, Cat no: PZ0198, Merck), all resuspended at 1 mg/mL in water, and DMPS (2,3-dimercapto-1-propanesulfonic acid sodium salt monohydrate, 95%, Cat no: H56578, Alfa Aesar) resuspended at 1mg/mL in PBS. Groups of five mice received experimental doses that consisted of either: (a) venom only (4 x LD_50_ dose) or (b) venom (4 x LD_50_ dose) with drug (118 μg) or (c) drug only (118 μg) to assess drug safety. The control group was the venom only group, against which all drug treatments were compared. Each experimental group comprised 5 animals as this was previously determined to be the minimum number of animals required to produce statistically significant results^39^. No randomisation was used to allocate experimental groups - mice were randomly allocated into cages of five prior to the experiment, and each cage formed one treatment group. No criteria for including or excluding animals was applied, and all data points were included in analyses. A total of 45 mice were used. All experimental doses were prepared to a volume of 200 μL in PBS and incubated at 37 °C for 30 mins prior to intravenous injection via the tail vein. Animals were monitored for humane endpoints (loss of righting reflex, seizure, external haemorrhage) for six hours, and any animals showing such signs were immediately euthanised by rising concentrations of carbon dioxide. All observations were performed by mixed gender experimenters who were blinded to the drug group allocation. Time of death, number of deaths and number of survivors were recorded, where deaths and times of death represent implementation of humane endpoint-dictated euthanasia. Kaplan-Meier survival plots were generated using GraphPad Prism 9.0 (GraphPad Software, San Diego, USA) and log-rank (Mantel-Cox) tests were used to statistically compare the survival times between groups treated with and without drug.

## 3. Results

### 3.1 Small molecule drugs have varying effects on the procoagulant activity of crude *D. typus* venom

The addition of *D. typus* venom to recalcified bovine plasma in the coagulation assay resulted in earlier stimulation of clotting compared to the no venom control (natural clotting), highlighting the procoagulant nature of this venom (Figure 1A). The effects of the small molecules against the procoagulant activity of crude *D. typus* venom are shown in Figure 1. Weak inhibitory effects were observed for the metal chelators dimercaprol and DMPS at micromolar concentrations. As shown in Figure 1B, concentrations of 160 μM showed strong inhibitory activity for both drugs, but this inhibitory effect rapidly decreased at lower concentrations, with no effect observed at concentrations of 10 μM and lower. IC_50_ values were determined to be 77.7 μM for dimercaprol and 120 μM for DMPS (Figure 1C), although due to the small number of data points between 0 and 100% inhibition these values must be interpreted with caution and 95% confidence intervals were unable to be calculated. Contrastingly, the peptidomimetic matrix metalloproteinase inhibitors marimastat and prinomastat potently neutralised the procoagulant effects of *D. typus* venom (Figure 1B). The IC_50_ values for marimastat and prinomastat were determined to be 34.2 nM (95% CI 24.2 to 48.5 nM) and 75.6 nM (95% CI 58.6 to 97.7 nM) respectively (Figure 1C), demonstrating that marimastat was significantly more potent in this assay than prinomastat (p = 0.01). The PLA_2_ inhibitor varespladib (control non-SVMP inhibiting drug used throughout) had no neutralising effect on venom-induced coagulation at any of the tested drug concentrations (Supplemental File S1), a result in line with our expectations of procoagulant venom activity being mediated by SVMP toxins.

**Figure 1.**
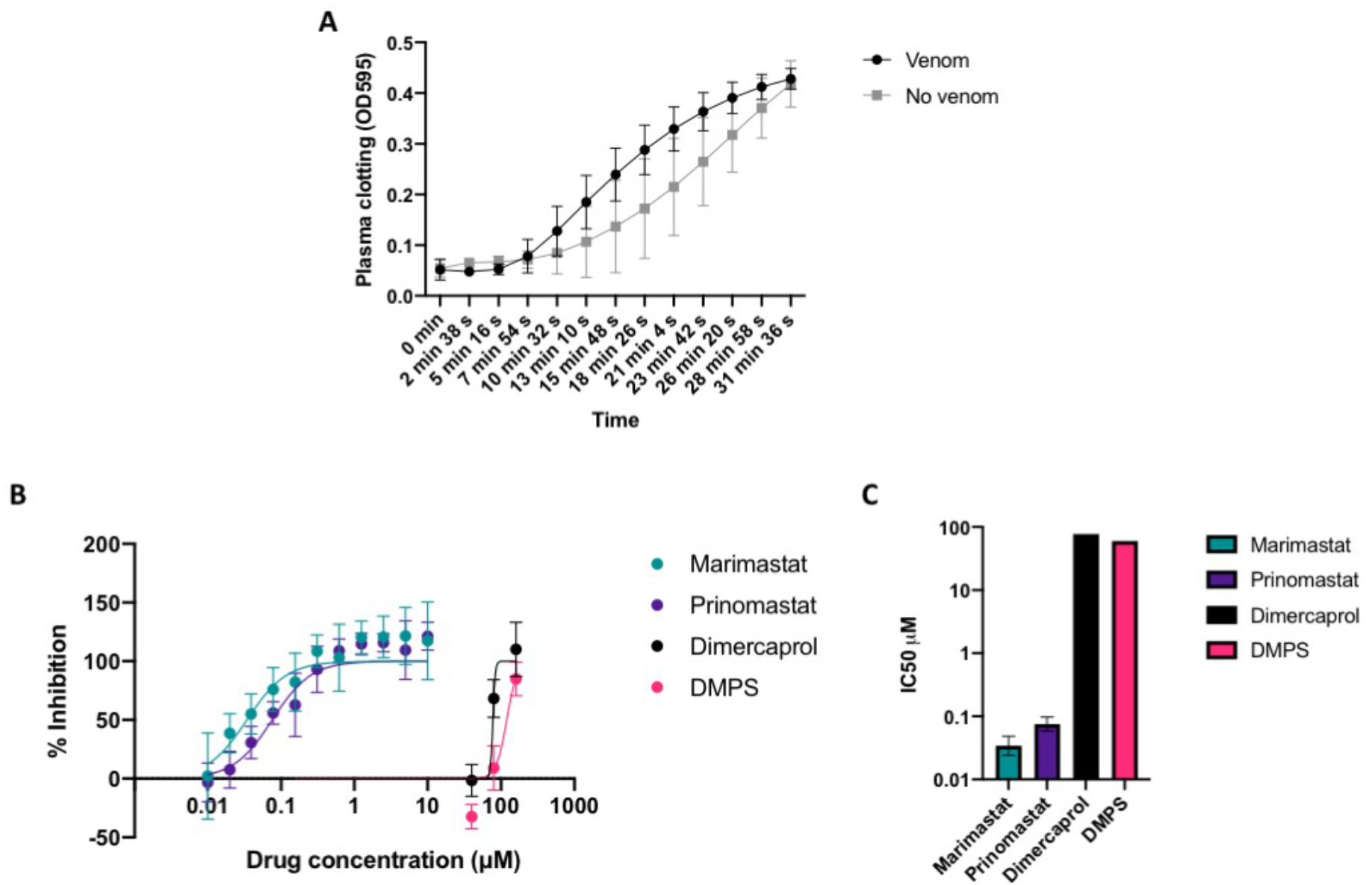
Effects of *D. typus* venom, and inhibition by small molecule drugs, on *in vitro* plasma clotting measured by absorbance at 595 nm (OD_595_). Inhibitory activity is expressed as a percentage of normal clotting, where 100% inhibition represents return of clotting to normal plasma clotting levels. **A)** Clotting as indicated by the increase in OD_595_ in the presence of crude *D. typus* venom (black circles) compared to normal clotting in the absence of venom (grey squares). Data points represent the mean of twelve individual values recorded over three independent technical replicates, and error bars represent standard deviation. **B)** Inhibitory activity of marimastat (teal circles), prinomastat (dark purple circles), dimercaprol (light purple circles) and DMPS (pink circles) over a two-fold serial dilution curve, from which IC_50_ values were calculated. Data points represent the mean of six individual values recorded over three independent technical replicates, and error bars represent standard deviation. **C)** Calculated IC_50_ values for marimastat, prinomastat, dimercaprol and DMPS. Data points represent the best fit IC_50_ value and error bars represent 95% confidence intervals (not calculated for dimercaprol and DMPS).

### 3.2 Inhibitory effects of small molecule drugs on nanofractionated *D. typus* venom coagulotoxins

To further characterise the coagulopathic activity of *D. typus* venom, and to better explore the specificity of the various small molecule inhibitors against specific toxins, we repeated the coagulation assay experiments using nanofractionated venom. As previously described, this method uses fractionated venom as the basis for measurements of the velocity of clotting in different wells in comparison to control wells, with procoagulant toxins producing positive peaks in the resulting bioassay chromatogram and anticoagulant toxins producing negative peaks^37^. In the venom-only analysis, broad positive peaks (18.4-22.0 min) were observed for both the ‘very fast coagulation’ chromatograms and the ‘slightly/medium increased coagulation’ chromatograms, indicative of an overall procoagulant effect of the venom. Detected bioactivities of *D. typus* venom were correlated with previously generated LC-MS and proteomics data^29^, and a candidate toxin mass of 23 kDa was identified for the pro-coagulant activity, which is within the range of SVMPs. The inhibitory effects of the matrix metalloproteinase inhibitors marimastat and prinomastat on nanofractionated *D. typus* venom toxins are depicted in Figure 2A and 2B respectively. The peaks in the very fast and slight/medium increased coagulation chromatograms decreased with increasing concentrations of both marimastat and prinomastat, indicative of a dose-dependent restoration of normal clotting velocity. All coagulopathic activities were inhibited at 0.8 μM marimastat and 0.16 μM prinomastat for very fast coagulation chromatograms, and at 4 μM for both inhibitory molecules for slightly/medium increased coagulation activity. In terms of the metal chelators dimercaprol and DMPS, their inhibitory effects on nanofractionated *D. typus* venom toxins are depicted in Figure 2C and 2D, respectively. By increasing the concentration range of dimercaprol, the procoagulation activity of *D. typus* venom was inhibited, with very fast coagulation activity fully inhibited at 4 μM, and slightly/medium increased coagulation activity at 20 μM. However, no substantial inhibition of procoagulant venom activity was observed with DMPS at any tested concentration up to 20 μM. These results reflect the considerable potency differences observed between the peptidomimetic inhibitors and the metal chelators in the crude venom plasma bioassays.

**Figure 2.**
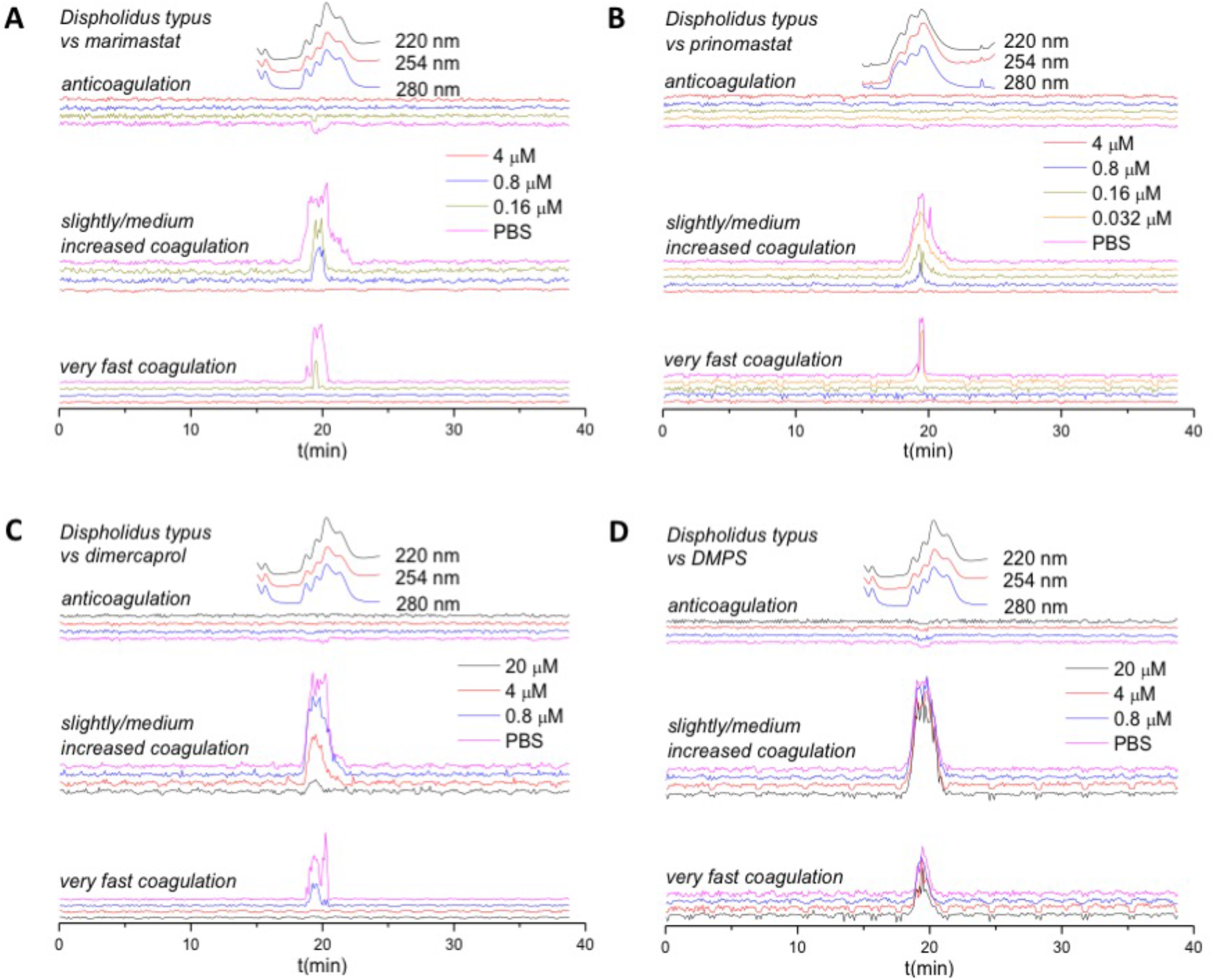
Reconstructed coagulation chromatograms for nanofractionated *D. typus* venom toxins in the presence of different concentrations of A) marimastat, B) prinomastat, C) dimercaprol, and D) DMPS. The negative peaks indicate anticoagulant activity where velocity is lower than the assay solution in control wells without venom toxins, and the positive peaks indicate pro-coagulant activity where velocity is higher than that in control wells without venom toxins. The top superimposed chromatograms are characteristic profiles of the UV trace at 220, 254 and 280 nm. PBS indicates venom only samples where PBS was used as a control for the inhibitors. Traces with different colours indicate different concentrations (final) of inhibitors in the assay.

A very weak signal (peak centre at 19.6 min) was also detected in terms of anticoagulant venom activity. A previous study observed a much clearer negative anticoagulant peak with *D. typus* venom, though the venom was applied at a five-fold higher concentration (5.0 mg/mL venom) than that used in this study (1.0 mg/mL venom)^29^. While varespladib has previously been demonstrated to be a potent inhibitor of anticoagulant venom activities induced by PLA_2_ toxins^33,40–42^, in this study it produced no inhibitory effects on venom-induced coagulation, whether procoagulant or anticoagulant, at the maximal drug dose tested (20 μM) (Supplemental File S2). However, due to the weak anticoagulant venom activity observed in these experiments, the assay window for measuring such inhibition is limited.

### 3.3 Small molecule drugs inhibit crude *D. typus* venom SVMP activity

To determine the specific inhibitory effects of the selected small molecule drugs on SVMP toxin activity, we performed IC_50_ screens of the five different toxin inhibitors in a previously defined kinetic enzymatic assay SVMP using crude *D. typus* venom. As previously observed^43^, *D. typus* venom demonstrated strong SVMP-specific activity in this assay. The inhibitory effects of the matrix metalloproteinase inhibitors marimastat and prinomastat and the metal chelators DMPS and dimercaprol against the venom SVMP activity of *D. typus* are displayed in Figure 3. The peptidomimetic inhibitors marimastat and prinomastat demonstrated nanomolar IC_50_ values of 14.5 (95% CI 13.9 to 15.2 nM) and 25.9 nM (95% CI 22.6 to 29.7 nM) respectively, and complete inhibition of venom activity at 156 nM for marimastat and 2.5 μM for prinomastat (Figure 3A). Inhibition of SVMP activity by dimercaprol and DMPS as measured by IC_50_ values was significantly lower than that observed for marimastat and prinomastat (p < 0.02 for all comparisons). Dimercaprol and DMPS demonstrated highly comparable inhibition of *D. typus* SVMP activity with complete inhibition obtained at 40 μM and IC_50_ values of 6.68 (95% CI 6.36 to 7.02 μM) and 6.61 μM (95% CI 6.38 to 6.87 μM), respectively (Figure 3B), with no significant differences between the two IC_50_ values. As anticipated, the control drug used in this study, the PLA_2_ inhibitor varespladib, showed no inhibitory activity at any of the concentrations tested (maximum concentration 10 μM) (Supplemental File S3).

**Figure 3.**
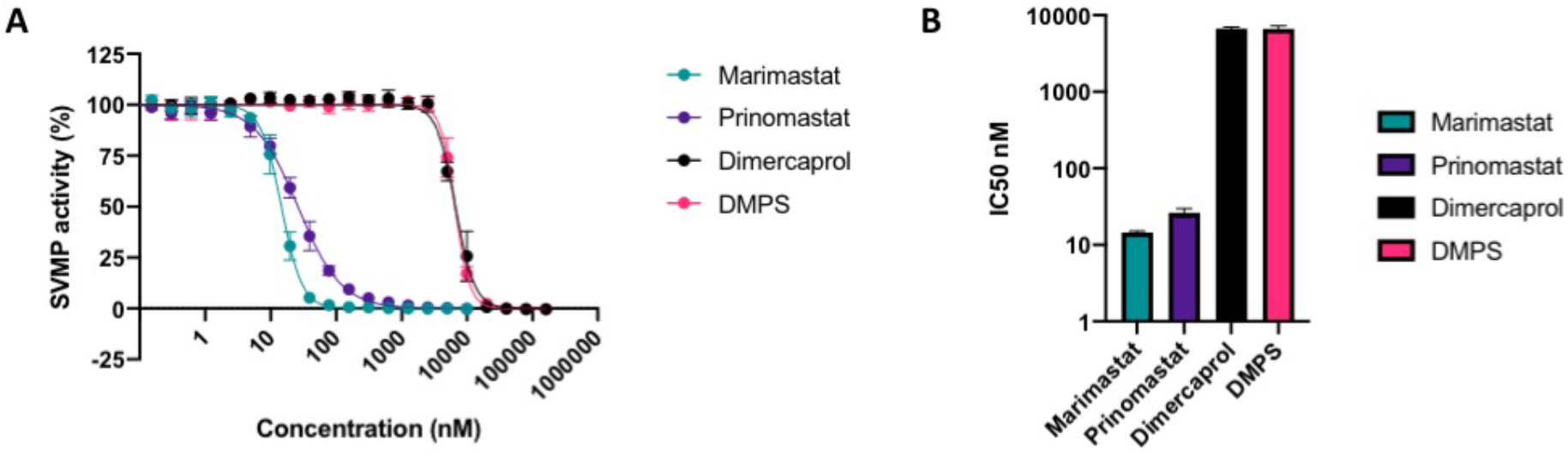
*In vitro* inhibition of *D. typus* crude venom SVMP activity by small molecule inhibitors. A) SVMP activity of *D. typus* crude venom in the presence of marimastat (teal circles), prinomastat (dark purple circles), dimercaprol (light purple circle) and DMPS (pink circles) over a two-fold serial dilution curve ranging from which IC_50_ values were calculated. Data points represent the percentage of crude venom SVMP activity generated from the mean of four individual values recorded over two independent technical replicates, and error bars represent standard deviation. **B)** IC_50_ values of SVMP inhibition for marimastat, prinomastat, dimercaprol and DMPS.

### 3.4 Marimastat and DMPS provide some protection against *D. typus* venom-induced lethality *in vivo*

Given that inhibition of SVMP and coagulotoxic activities *in vitro* have previously been demonstrated to translate into varying degrees of *in vivo* protection against systemic envenoming^26,36^ we next tested the capability of the four SVMP-inhibiting small molecule drugs to protect against *D. typus* venom-induced lethality *in vivo*. To do so, we used a modified version of the WHO-recommended protocol of murine venom neutralisation (ED_50_ assay). All five experimental animals treated intravenously with 4 x LD_50_ doses of *D. typus* venom (90 μg) succumbed to the lethal venom effects within the first hour of the experiment (mean 17 minutes, range 1 to 40 minutes), as shown in Figure 4. The intravenous co-delivery of preincubated drugs with *D. typus* venom revealed that both prinomastat and dimercaprol failed to protect against venom induced lethality at the single therapeutic dose tested (118 μg), with all experimental animals in these groups succumbing to venom lethality within the first 30 minutes (mean 7.8 minutes for both groups; prinomastat range 4 to 21 minutes; dimercaprol range 1 to 18 minutes), in a highly comparable manner to the venom only control. Contrastingly, DMPS and marimastat both showed a significant degree of protection against *D. typus* venom-induced lethality. Three of the five experimental animals dosed with DMPS were protected for the duration of the experiment, with two deaths occurring within the first hour, resulting in a mean survival time of 224 minutes compared to 17.2 minutes in the venom only group (log-rank test, p = 0.047). Of the animals dosed with marimastat, four of the five animals were protected for the duration of the experiment, with mean survival times of 316.2 minutes compared to 17.2 minutes in the venom only group (log-rank test, p = 0.002). The single non-surviving experimental animal in this drug group was euthanised at 141 minutes. No adverse effects were observed in experimental animals dosed with any of the drugs only controls and, consequently, all survived the duration of the experiment (Figure 4).

**Figure 4.**
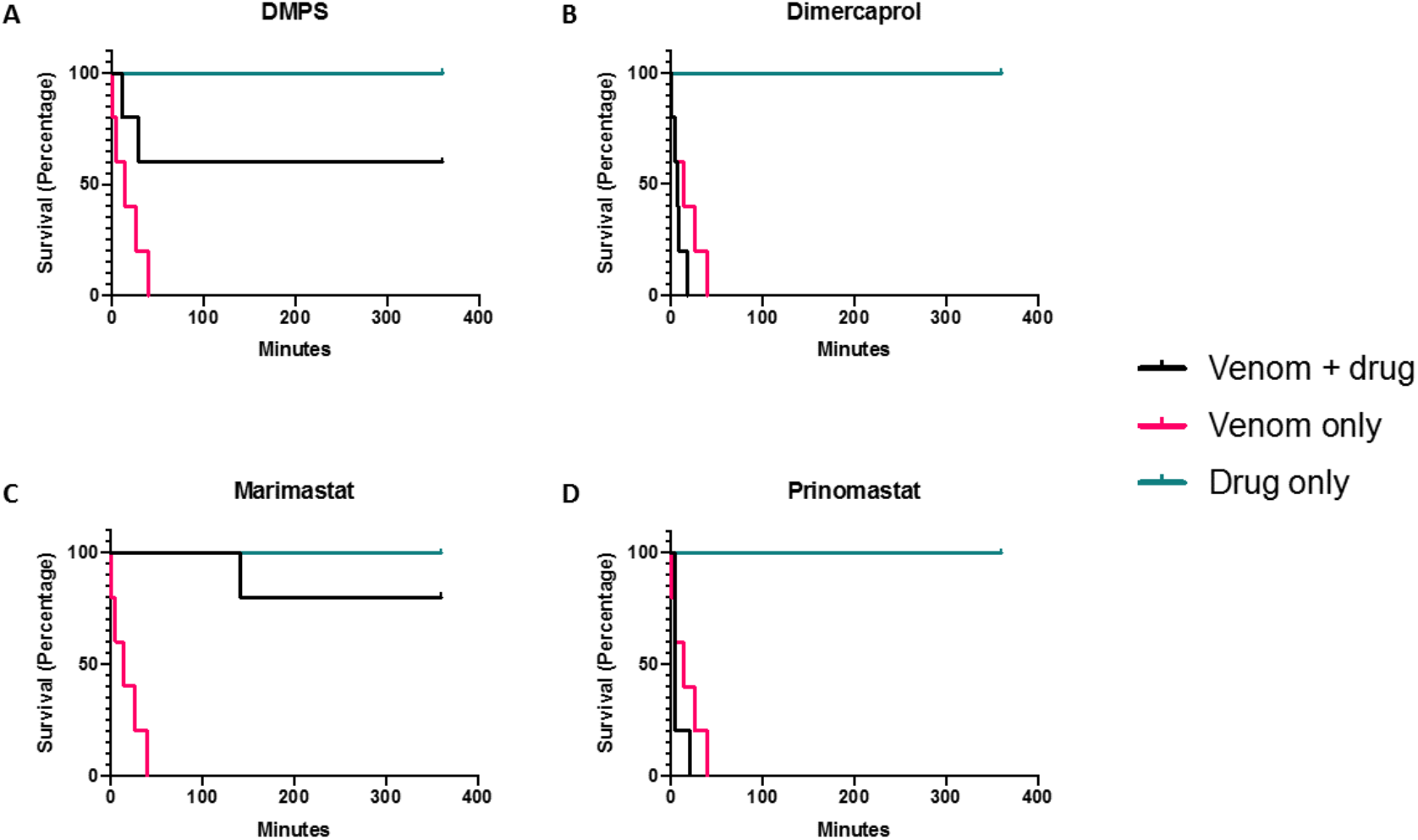
The small molecule drugs marimastat and DMPS significantly increase the survival times of mice receiving lethal doses of *D. typus* venom. The data is shown in Kaplan-Meier survival graphs for experimental animals (n=5 per group, except prinomastat only group where n=1) treated with either: 90 μg (4 x LD_50_) of *D. typus* venom only (magenta), 118 μg of drug only (cyan) or 90 μg venom and 118 μg drug (black). Treatments were pre-incubated at 37 °C for 30 minutes prior to intravenous injection via the tail vein and animals were monitored for 6 hours. Data is shown for: **A)** DMPS, **B)** dimercaprol, **C)** marimastat, and **D)** prinomastat

## 4. Discussion

Conventional animal-derived antivenom, although a life-saving treatment, has numerous deficiencies that impair its utility in the treatment of snakebite. Although a seemingly effective antivenom for treating *D. typus* envenomation is available in South Africa, it is often difficult to source in other regions of the continent and can be catastrophically unaffordable for patients^13^, and this scenario encapsulates the challenges faced by snakebite victims world over. There is therefore an urgent need to develop alternative/supplementary therapeutics that are stable, effective, affordable and available in remote rural areas where medical access is limited. Small molecule inhibitors that can broadly neutralise a class of key toxins in snake venom following oral administration are possible solutions in this regard^20^ and Phase II clinical trials for small molecule inhibitors of snakebite are underway^44^. Rapid-onset pathologies such as VICC, together with tissue damage induced by SVMPs, are only partially neutralised by antibody based antivenoms^45^, which suffer from poor tissue distribution due to the inherent large size of antibodies^46^. By contrast, the drastically smaller size of the inhibitors tested in this study enables favourable properties of rapid and effective tissue penetration and potential for oral delivery, due to their pharmacokinetic and physicochemical properties^34,35,47^. Moreover, repurposing these small molecule inhibitors that are either licensed drugs or phase I-approved drug candidates could significantly shorten drug development times as safety profiles, pharmacokinetics, bioavailability and tolerance data on these molecules have already obtained^47–49^. Current evidence of the utility of small molecule inhibitors against snakebite indicates that they may be particularly effective as first line, early intervention, therapeutics and/or bridging therapies for initial and adjunct treatment in community settings, before patients are able to access antivenom in healthcare centres^18,28,48^.

In this study we assessed the ability of four small molecule inhibitors to neutralise *D. typus* venom toxin activities *in vitro* and *in vivo*. In line with previous findings^8–11^, our study demonstrates that *D. typus* venom toxicity is largely conferred by SVMP toxins, which are likely responsible for causing coagulopathy *in vivo*. Our data from *in vitro* assays of *D. typus* venom activity demonstrate that the matrix metalloproteinase inhibitors marimastat and prinomastat are potent inhibitors of the SVMP-mediated procoagulant effects of this venom, with both compounds demonstrating similar inhibitory activity in *in vitro* assays of plasma coagulation and SVMP activity. Indeed, marimastat and prinomastat showed nanomolar IC_50_ values in the crude venom plasma coagulation assay and crude venom SVMP assay, and both drugs showed inhibitory effects at low micromolar concentrations in the plasma coagulation assay with nanofractionated venom. However, and in contrast with *in vitro* SVMP-inhibiting prowess, in the *in vivo* assays, all animals in the prinomastat group succumbed to lethality from *D. typus* venom in murine models of envenomation, whilst an equivalent amount of marimastat conferred 80% protection. This was unexpected and hints at different levels of drug exposure and metabolism in this single dose intravenous-delivered model, as both matrix metalloproteinase inhibitors showed potent inhibitory activity in the *in vitro* assays and both have been shown in other studies to neutralise the *in vivo* lethal effects of *E. ocellatus* in murine models^24,26^. The metal chelators dimercaprol and DMPS demonstrated lower potency than marimastat and prinomastat in the *in vitro* studies of SVMP activity and plasma coagulation, whilst the PLA_2_ inhibitor varespladib, used as a non-SVMP inhibiting control, produced no inhibitory effects on venom. Of the two metal chelators, DMPS showed the weakest inhibitory activity in *in vitro* assays of venom bioactivity. Despite this reduced *in vitro* potency in comparison with the peptidomimetic matrix metalloproteinase inhibitors, DMPS conferred a degree of protection against venom lethality in murine models of envenomation, with significant increases in mean survival times and 60% of mice surviving until the end of the experimental time window. None of the experimental animals dosed with the other metal chelator, dimercaprol, survived the experiment, despite the comparable mechanism of action and *in vitro* inhibitory potency in the coagulation and SVMP assays. These results contrast with our previous preclinical study investigating *E. ocellatus* envenoming, which found that both dimercaprol and DMPS provided protection against lethal effects in this same intravenous murine model, though are consistent with DMPS exhibiting superior preclinical efficacy^26^.

The notable discrepancies between the *in vitro* and *in vivo* experiments described herein exemplifies the complexity associated with relying *on in* vitro potency-based screens as a means to predict the efficacy of small molecule drugs in *in vivo* experiments. While efficacy data gained from *in vitro* experiments is undoubtedly an essential prerequisite prior to preclinical efficacy testing, substantial differences in inhibitor potency at this step does not preclude preclinical efficacy, which ultimately is dictated by drug exposure. Equally, the preclinical model utilised here, consisting of the pre-incubation of drug with venom followed by codelivery intravenously, is largely detached from the clinical scenario of a snakebite. While this is the WHO-recommended method for preclinical assessment of antivenom efficacy, and thus is a logical starting point for assessing preclinical efficacy, this method does not reflect the biodistribution of venom during early envenomation, uses a non-clinically relevant route of venom injection, and does not take into account the pharmacokinetics/pharmacodynamics of the unbound test inhibitor^50^. Thus, the lack of efficacy observed with prinomastat (compared with marimastat) here, for example, may be the result of a lack of dose optimisation and thus sub-optimal exposure. Future work is required to better define the pharmacokinetic and pharmacodynamic profiles of small molecule drugs in preclinical models of snakebite envenoming to inform the design of optimised pre-clinical dosing regimens applicable for use in more biologically realistic models of envenoming (e.g. “challenge then treat models”)^26,34,36,46,51^.

In sub-Saharan Africa, VICC is only known to be commonly caused by *Echis* spp. and *D. typus,* and the venoms of these snakes have been shown to converge on similar SVMP-rich venom composition profiles^9^, suggesting that *D. typus* venom may be amenable to neutralisation by previously identified inhibitors of *Echis* venoms. This study investigated the ability of repurposed small molecule inhibitors to effectively neutralise *D. typus* venom activity *in vitro* and *in vivo,* and identified the SVMP inhibiting drugs DMPS and marimastat as two lead compounds that provide a significant degree of preclinical protection against the lethal effects of *D. typus* venom. Previous studies have demonstrated that both DMPS and marimastat also provide preclinical efficacy against *E. ocellatus* envenoming^26,36^, and this study therefore expands the range of snake species that victims of envenoming could potentially benefit from receiving an early intervention (e.g. oral) small molecule therapeutic. These findings therefore provide a strong rationale for the future clinical evaluation of the efficacy of such small molecule drugs in all cases of diagnostically indicated VICC following snakebite envenoming in sub-Saharan Africa.

## Acknowledgements

We thank the staff in the Biomedical Service Unit of the University of Liverpool for their support in the maintenance and care of the study mice.

## Author contributions

Conceptualization – JK, NRC

Methodology – SKM, RHC, CX

Investigation – SKM, RHC, CX, AW, SRH, RJE, JA, EC, RAH, NRC,

Data curation – SKM, RHC, CX, AW

Formal analysis – SKM, RHC, CX, AW

Original draft preparation – SKM, RHC, CX, NRC, JK

Editing - all authors

## References

1. Halilu S, Iliyasu G, Hamza M, Chippaux JP, Kuznik A, Habib AG. Snakebite burden in Sub-Saharan Africa: estimates from 41 countries. Toxicon. 2019;159:1–4. doi:10.1016/j.toxicon.2018.12.002

2. Longbottom J, Shearer FM, Devine M, et al. Vulnerability to snakebite envenoming: a global mapping of hotspots. The Lancet. 2018;392(10148):673–684. doi:10.1016/S0140-6736(18)31224-8

3. Harrison RA, Casewell NR, Ainsworth SA, Lalloo DG. The time is now: a call for action to translate recent momentum on tackling tropical snakebite into sustained benefit for victims. Trans R Soc Trop Med Hyg. 2019;113(12):835–838. doi:10.1093/trstmh/try134

4. Berling I, Isbister GK. Hematologic Effects and Complications of Snake Envenoming. Transfus Med Rev. 2015;29(2):82–89. doi:10.1016/j.tmrv.2014.09.005

5. Lakier JB, Fritz VU. Consumptive coagulopathy caused by a boomslang bite. South Afr Med J Suid-Afr Tydskr Vir Geneeskd. 1969;43(34):1052–1055.

6. Matell G, Nyman D, Werner B, Wilhelmsson S. Consumption coagulopathy caused by a boomslang bite: A case report. Thromb Res. 1973;3(2):173–182. doi:10.1016/0049-3848(73)90067-4

7. Gomperts ED, Demetriou D. Laboratory studies and clinical features in a case of boomslang envenomation. South Afr Med J Suid-Afr Tydskr Vir Geneeskd. 1977;51(6):173–175.

8. Debono J, Dobson J, Casewell NR, et al. Coagulating Colubrids: Evolutionary, Pathophysiological and Biodiscovery Implications of Venom Variations between Boomslang (Dispholidus typus) and Twig Snake (Thelotornis mossambicanus). Toxins. 2017;9(5):171. doi:10.3390/toxins9050171

9. Ainsworth S, Slagboom J, Alomran N, et al. The paraspecific neutralisation of snake venom induced coagulopathy by antivenoms. Commun Biol. 2018;1(1):1–14. doi:10.1038/s42003-018-0039-1

10. Debono J, Dashevsky D, Nouwens A, Fry BG. The sweet side of venom: Glycosylated prothrombin activating metalloproteases from Dispholidus typus (boomslang) and Thelotornis mossambicanus (twig snake). Comp Biochem Physiol Part C Toxicol Pharmacol. 2020;227:108625. doi:10.1016/j.cbpc.2019.108625

11. Pla D, Sanz L, Whiteley G, et al. What killed Karl Patterson Schmidt? Combined venom gland transcriptomic, venomic and antivenomic analysis of the South African green tree snake (the boomslang), Dispholidus typus. Biochim Biophys Acta Gen Subj. 2017;1861(4):814–823. doi:10.1016/j.bbagen.2017.01.020

12. Kamiguti AS, Theakston RDG, Sherman N, Fox JW. Mass spectrophotometric evidence for P-III/P-IV metalloproteinases in the venom of the Boomslang (Dispholidus typus). Toxicon. 2000;38(11):1613–1620. doi:10.1016/S0041-0101(00)00089-1

13. Krüger HJ, Lemke FG. Fatal Boomslang bite in the Northern Cape. Afr J Emerg Med. 2019;9(1):53–55. doi:10.1016/j.afjem.2018.12.006

14. de Silva HA, Ryan NM, de Silva HJ. Adverse reactions to snake antivenom, and their prevention and treatment. Br J Clin Pharmacol. 2016;81(3):446–452. doi:10.1111/bcp.12739

15. Potet J, Smith J, McIver L. Reviewing evidence of the clinical effectiveness of commercially available antivenoms in sub-Saharan Africa identifies the need for a multi-centre, multi-antivenom clinical trial. PLoS Negl Trop Dis. 2019;13(6):e0007551. doi:10.1371/journal.pntd.0007551

16. Casewell NR, Cook DAN, Wagstaff SC, et al. Pre-Clinical Assays Predict Pan-African Echis Viper Efficacy for a Species-Specific Antivenom. PLoS Negl Trop Dis. 2010;4(10):e851. doi:10.1371/journal.pntd.0000851

17. World Health Organization. Guidelines for the Prevention and Clinical Management of Snakebite in Africa.; 2010. Accessed December 21, 2021. https://www.who.int/publications-detail-redirect/9789290231684

18. Bulfone TC, Samuel SP, Bickler PE, Lewin MR. Developing Small Molecule Therapeutics for the Initial and Adjunctive Treatment of Snakebite. J Trop Med. 2018;2018:1–10. doi:10.1155/2018/4320175

19. Williams HF, Layfield HJ, Vallance T, et al. The Urgent Need to Develop Novel Strategies for the Diagnosis and Treatment of Snakebites. Toxins. 2019;11(6). doi:10.3390/toxins11060363

20. Clare RH, Hall SR, Patel RN, Casewell NR. Small Molecule Drug Discovery for Neglected Tropical Snakebite. Trends Pharmacol Sci. 2021;42(5):340–353. doi:10.1016/j.tips.2021.02.005

21. Rucavado A, Escalante T, Franceschi A, et al. Inhibition of local hemorrhage and dermonecrosis induced by Bothrops asper snake venom: Effectiveness of early in situ administration of the peptidomimetic metalloproteinase inhibitor batimastat and the chelating agent CaNa EDTA. Am J Trop Med Hyg 63. Published online 2000.

22. Layfield HJ, Williams HF, Ravishankar D, et al. Repurposing Cancer Drugs Batimastat and Marimastat to Inhibit the Activity of a Group I Metalloprotease from the Venom of the Western Diamondback Rattlesnake, Crotalus atrox. Toxins. 2020;12(5):309. doi:10.3390/toxins12050309

23. Warrell, D.A., Davidson N, Greenwood B, et al. Poisoning by bites of the saw-scaled or carpet viper (Echis carinatus) in Nigeria. Q J Med. 1977;46:33–62.

24. Arias AS, Rucavado A, Gutiérrez JM. Peptidomimetic hydroxamate metalloproteinase inhibitors abrogate local and systemic toxicity induced by Echis ocellatus (saw-scaled) snake venom. Toxicon Off J Int Soc Toxinology. 2017;132:40–49. doi:10.1016/j.toxicon.2017.04.001

25. Xie C, Albulescu LO, Bittenbinder MA, et al. Neutralizing Effects of Small Molecule Inhibitors and Metal Chelators on Coagulopathic Viperinae Snake Venom Toxins. Biomedicines. 2020;8(9):297. doi:10.3390/biomedicines8090297

26. Albulescu LO, Hale MS, Ainsworth S, et al. Preclinical validation of a repurposed metal chelator as an early-intervention therapeutic for hemotoxic snakebite. Sci Transl Med. 2020;12(542). doi:10.1126/scitranslmed.aay8314

27. Xie C, Slagboom J, Albulescu LO, et al. Neutralising effects of small molecule toxin inhibitors on nanofractionated coagulopathic Crotalinae snake venoms. Acta Pharm Sin B. 2020;10(10):1835–1845. doi:10.1016/j.apsb.2020.09.005

28. Howes JM, Theakston RDG, Laing GD. Neutralization of the haemorrhagic activities of viperine snake venoms and venom metalloproteinases using synthetic peptide inhibitors and chelators. Toxicon Off J Int Soc Toxinology. 2007;49(5):734–739. doi:10.1016/j.toxicon.2006.11.020

29. Slagboom J, Mladić M, Xie C, et al. High throughput screening and identification of coagulopathic snake venom proteins and peptides using nanofractionation and proteomics approaches. PLoS Negl Trop Dis. 2020;14(4):e0007802. doi:10.1371/journal.pntd.0007802

30. Xie C, Albulescu LO, Still KBM, et al. Varespladib Inhibits the Phospholipase A2 and Coagulopathic Activities of Venom Components from Hemotoxic Snakes. Biomedicines. 2020;8(6):165. doi:10.3390/biomedicines8060165

31. Wang Y, Zhang J, Zhang D, Xiao H, Xiong S, Huang C. Exploration of the Inhibitory Potential of Varespladib for Snakebite Envenomation. Molecules. 2018;23(2):391. doi:10.3390/molecules23020391

32. Bryan-Quirós W, Fernández J, Gutiérrez JM, Lewin MR, Lomonte B. Neutralizing properties of LY315920 toward snake venom group I and II myotoxic phospholipases A2. Toxicon. 2019;157:1–7. doi:10.1016/j.toxicon.2018.11.292

33. Bittenbinder MA, Zdenek CN, Op den Brouw B, et al. Coagulotoxic Cobras: Clinical Implications of Strong Anticoagulant Actions of African Spitting Naja Venoms That Are Not Neutralised by Antivenom but Are by LY315920 (Varespladib). Toxins. 2018;10(12):516. doi:10.3390/toxins10120516

34. Lewin MR, Gutiérrez JM, Samuel SP, et al. Delayed Oral LY333013 Rescues Mice from Highly Neurotoxic, Lethal Doses of Papuan Taipan (Oxyuranus scutellatus) Venom. Toxins. 2018;10(10):380. doi:10.3390/toxins10100380

35. Lewin MR, Gilliam LL, Gilliam J, et al. Delayed LY333013 (Oral) and LY315920 (Intravenous) Reverse Severe Neurotoxicity and Rescue Juvenile Pigs from Lethal Doses of Micrurus fulvius (Eastern Coral Snake) Venom. Toxins. 2018;10(11). doi:10.3390/toxins10110479

36. Albulescu LO, Xie C, Ainsworth S, et al. A therapeutic combination of two small molecule toxin inhibitors provides broad preclinical efficacy against viper snakebite. Nat Commun. 2020;11(1):6094. doi:10.1038/s41467-020-19981-6

37. Still KBM, Nandlal RSS, Slagboom J, Somsen GW, Casewell NR, Kool J. Multipurpose HTS Coagulation Analysis: Assay Development and Assessment of Coagulopathic Snake Venoms. Toxins. 2017;9(12). doi:10.3390/toxins9120382

38. Zietek BM, Mayar M, Slagboom J, et al. Liquid chromatographic nanofractionation with parallel mass spectrometric detection for the screening of plasmin inhibitors and (metallo)proteinases in snake venoms. Anal Bioanal Chem. 2018;410(23):5751–5763. doi:10.1007/s00216-018-1253-x

39. World Health Organization. WHO Guidelines for the Production, Control and Regulation of Snake Antivenom Immunoglobulins. Geneva: World Health Organization.; 2018. https://www.who.int/snakebites/resources/Snake_antivenom_immunoglobulins_WHO_TRS1004_Annex5.pdf?ua=1

40. Kazandjian TD, Arrahman A, Still KBM, et al. Anticoagulant Activity of Naja nigricollis Venom Is Mediated by Phospholipase A2 Toxins and Inhibited by Varespladib. Toxins. 2021;13(5):302. doi:10.3390/toxins13050302

41. Youngman NJ, Walker A, Naude A, Coster K, Sundman E, Fry BG. Varespladib (LY315920) neutralises phospholipase A2 mediated prothrombinase-inhibition induced by Bitis snake venoms. Comp Biochem Physiol Part C Toxicol Pharmacol. 2020;236:108818. doi:10.1016/j.cbpc.2020.108818

42. Liu CC, Wu CJ, Hsiao YC, et al. Snake venom proteome of Protobothrops mucrosquamatus in Taiwan: Delaying venom-induced lethality in a rodent model by inhibition of phospholipase A2 activity with varespladib. J Proteomics. 2021;234:104084. doi:10.1016/j.jprot.2020.104084

43. Alomran N, Alsolaiss J, Albulescu LO, et al. Pathology-specific experimental antivenoms for haemotoxic snakebite: The impact of immunogen diversity on the in vitro cross-reactivity and in vivo neutralisation of geographically diverse snake venoms. PLoS Negl Trop Dis. 2021;15(8):e0009659. doi:10.1371/journal.pntd.0009659

44. Randomized, Double-Blinded, Placebo-Controlled Study to Evaluate the Safety, Tolerability, and Efficacy of a Multi-Dose Regimen of Oral Varespladib-Methyl in Subjects Bitten by Venomous Snakes. clinicaltrials.gov; 2021. Accessed November 15, 2021. https://clinicaltrials.gov/ct2/show/NCT04996264

45. Gutiérrez JM, Theakston RDG, Warrell DA. Confronting the Neglected Problem of Snake Bite Envenoming: The Need for a Global Partnership. PLOS Med. 2006;3(6):e150. doi:10.1371/journal.pmed.0030150

46. Gutiérrez JM, Solano G, Pla D, et al. Preclinical Evaluation of the Efficacy of Antivenoms for Snakebite Envenoming: State-of-the-Art and Challenges Ahead. Toxins. 2017;9(5):163. doi:10.3390/toxins9050163

47. Kini RM, Sidhu SS, Laustsen AH. Biosynthetic Oligoclonal Antivenom (BOA) for Snakebite and Next-Generation Treatments for Snakebite Victims. Toxins. 2018;10(12):534. doi:10.3390/toxins10120534

48. Lewin M, Samuel S, Merkel J, Bickler P. Varespladib (LY315920) Appears to Be a Potent, Broad-Spectrum, Inhibitor of Snake Venom Phospholipase A2 and a Possible Pre-Referral Treatment for Envenomation. Toxins. 2016;8(9):248. doi:10.3390/toxins8090248

49. Knudsen C, Ledsgaard L, Dehli RI, Ahmadi S, Sørensen CV, Laustsen AH. Engineering and design considerations for next-generation snakebite antivenoms. Toxicon Off J Int Soc Toxinology. 2019;167:67–75. doi:10.1016/j.toxicon.2019.06.005

50. Gutiérrez JM, Albulescu LO, Clare RH, et al. The Search for Natural and Synthetic Inhibitors That Would Complement Antivenoms as Therapeutics for Snakebite Envenoming. Toxins. 2021;13(7):451. doi:10.3390/toxins13070451

51. Knudsen C, Casewell NR, Lomonte B, Gutiérrez JM, Vaiyapuri S, Laustsen AH. Novel Snakebite Therapeutics Must Be Tested in Appropriate Rescue Models to Robustly Assess Their Preclinical Efficacy. Toxins. 2020;12(9):528. doi:10.3390/toxins12090528

